# Statistical context dictates the relationship between feedback-related EEG signals and learning

**DOI:** 10.1101/581744

**Authors:** Matthew R. Nassar, Rasmus Bruckner, Michael J. Frank

**Affiliations:** Robert J. & Nancy D. Carney Institute for Brain Science, Brown University, Providence RI 02912-1821, USA; Department of Neuroscience, Brown University, Providence RI 02912-1821, USA; Department of Cognitive, Linguistic, and Psychological Sciences, Brown University, Providence RI 02912-1821, USA; Department of Education and Psychology, Freie Universität Berlin, 14195 Berlin, Germany; Center for Lifespan Psychology, Max Planck Institute for Human Development, 14195 Berlin, Germany; International Max Planck Research School on the Life Course (LIFE), Berlin, Germany

## Abstract

Successful decision-making requires learning expectations based on experienced outcomes. This learning should be calibrated according to the surprise associated with an outcome, but also to the statistical context dictating the most likely source of surprise. For example, when occasional changepoints are expected, surprising outcomes should be weighted heavily, demanding increased learning. In contrast, when signal corruption is expected to occur occasionally, surprising outcomes can suggest a corrupt signal that should be ignored by learning systems. Here we dissociate surprising outcomes from the degree to which they demand learning using a predictive inference task and computational modeling. We show that the updating P300, a stimulus-locked electrophysiological response previously associated with adjustments in learning behavior, does so conditionally on the source of surprise. Larger P300 signals predicted greater learning in a changing context, but predicted less learning in a context where surprise was indicative of a one-off outlier (oddball). The conditional predictive relationship between the P300 and learning behavior was persistent even after adjusting for known sources of learning rate variability. Our results suggest that the P300 provides a surprise signal that is interpreted by downstream learning processes differentially according to statistical context in order to appropriately calibrate learning across complex environments.

## Introduction

People are capable of rationally adjusting the degree to which they incorporate new information into their beliefs about the world (1–5). In environments that include discontinuous changes (changepoints) normative learning requires increasing learning when beliefs are uncertain or when observations are most surprising (2,6). Human participants display both of these tendencies, albeit to varying degrees (2,6,7).

A major open question in the learning domain is how the brain achieves such apparent adjustments in learning rate. This question has fueled a number of recent studies that have identified neural correlates of surprise in functional magnetic resonance imaging (fMRI) (8), electroencephalography (EEG) (9,10), and pupil signals (6) that predict subsequent learning behavior. These signals might reflect candidate mechanisms for a general system to adjust learning rate (1,11,12), yet the generality has yet to be established outside of discontinuously changing environments, where surprise and learning are tightly coupled.

The relationship between surprise and learning is complex and depends critically on the overarching statistical context. We refer to learning as the degree to which an observed prediction error promotes measurable behavioral updating. While changing environments require increased learning in the face of surprising information, stable environments with outliers (“oddballs”), dictate less learning from surprising information (4). People are capable of this type of robust learning rate adjustment that deemphasizes surprising information (3,4,13), yet the learning signals measured under such conditions do not correspond directly to those observed in changing environments. Most notably, a number of candidate learning signals measured through fMRI do not reflect learning rate when considering a broader set of statistical contexts (4).

However, prior studies on EEG correlates of learning seem to favor the idea that a late, stimulus-locked positivity referred to as the P300, tracks learning in a broader range of statistical contexts. The central parietal component of the P300 (P3b) reflects surprise (14) and relates to learning (15) even after controlling for the degree of surprise in changing environments (9,10). In a stationary environment where integration of sequential samples is required to make a subsequent decision, a late posterior positivity, reminiscent of the P300, predicts the degree to which a particular sample influences the subsequent decision (16). Interestingly, within this particular task, more surprising outcomes tended to exert less influence on decisions (3,13), suggesting that this late positivity might provide a general learning or updating signal, irrespective of statistical context. This idea would be in line with a prominent theory of P3b function, which emphasizes its role in updating context representations – sometimes defined in terms of items stored in working memory (17–20).

Here we tested the idea that the P3b provides a general learning signal that is independent of the statistical context. In particular, we measured learning behavior using a modified predictive inference task and normative learning model and examined how learning behavior and surprise related to evoked potentials measured through EEG. We found that people are capable of contextually adjusting learning in response to surprise: they tended to learn more from surprising outcomes when those outcomes were indicative of changepoints, but learned less from surprising outcomes when those outcomes were indicative of an oddball. Outcome evoked potentials reminiscent of a parietal P300 were related to surprising events irrespective of context. The magnitude of this P300 response on a given trial positively predicted learning in the presence of changepoints, but negatively predicted learning in the presence of oddballs. These relationships persisted even when controlling for variability in learning behavior that could be explained by the best behavioral model. Taken together these findings suggest that the P300 does not naively reflect increased behavioral updating, but may play a role in adaptively increasing or decreasing learning in response to surprising information, depending on the statistical context.

## Results

We used EEG to measure electrophysiological signatures of feedback processing while participants performed a modified predictive inference task (2) designed to dissociate surprise from learning. Predictions were made in the context of a video game that required participants to place a shield at a location on a circle in order to block cannonballs that would be fired from a cannon located at the center of the circle (Fig 1A). Surprise and learning were manipulated independently using two different task conditions. In the *oddball* condition, the aim of the cannon drifted slowly from one trial to the next (Fig 1B, dotted line) and cannonball locations were distributed around the point of cannon aim (Fig 1B, green points nearby dotted line) or, occasionally and unpredictably, uniformly distributed around the circle (*oddballs*; see green point on trial 11 of Fig 1B for example). In the *changepoint* condition, the cannon aim remained constant for an unpredictable duration, and was then re-aimed at a new location on the circle at random (*changepoints*; Fig 1C, dotted line). Cannonball locations were always distributed around the point of cannon aim in this condition (Fig 1C, green points).

**Figure 1:**
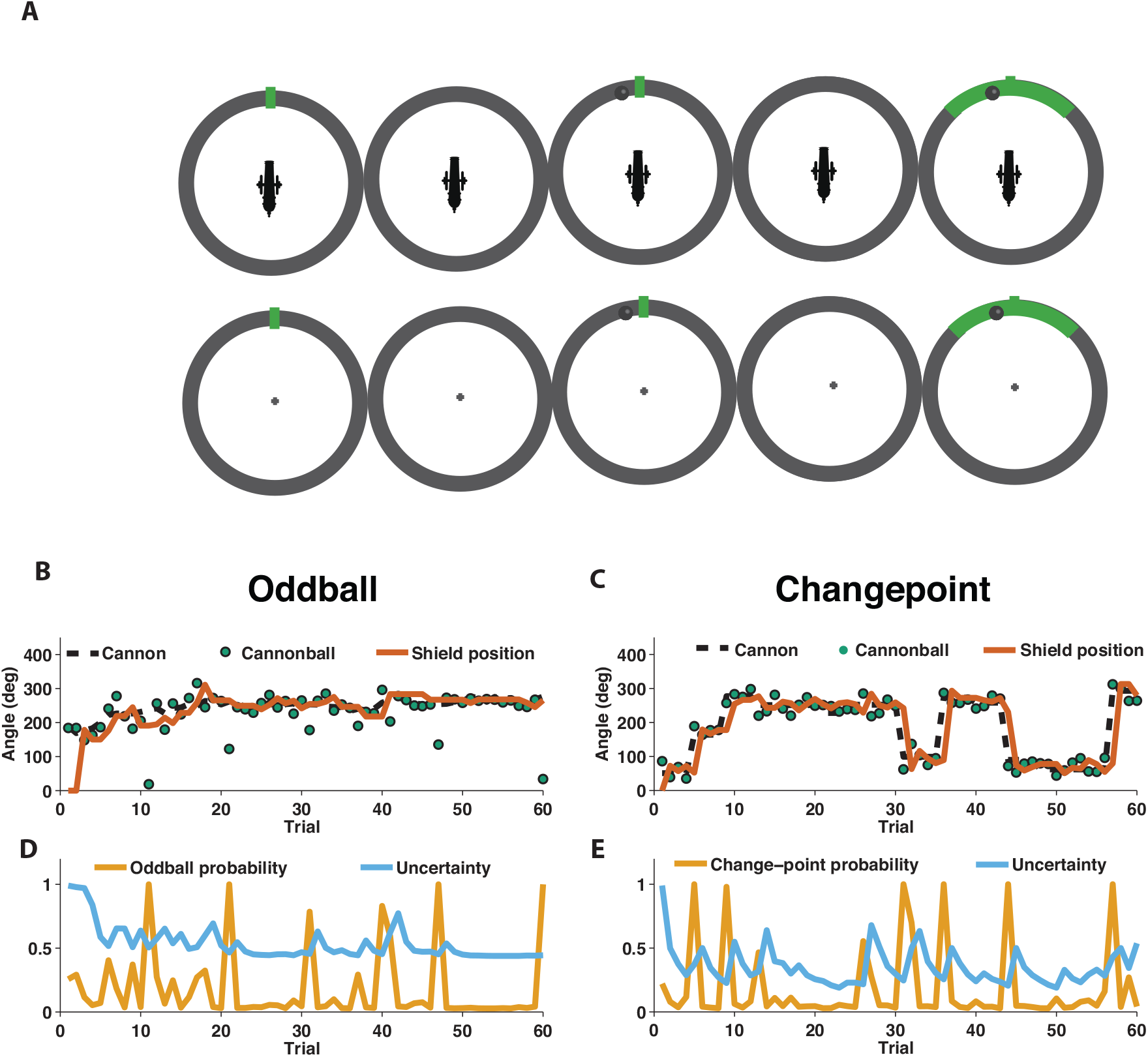
Measuring learning in different statistical contexts with a predictive inference task. **A)** Participants were trained to place the center of a shield (green tick; prediction phase [left]) at the aim location of a cannon (training task; top) in order to block a cannonball shot from it (outcome phase [top middle]) with a shield that varied in size from trial to trial and was revealed at the end of the trial (shield phase [top right]). After training, participants were asked to complete the same task, but without a visual depiction of the cannon, which required them to infer the aim of the cannon based on the sequence of previously observed cannonballs (test task; [bottom]). **B)** In oddball blocks, cannon aim (dotted black line) followed a random walk and cannonball locations were typically drawn from a Von Mises distribution centered on the true cannon aim (green points), but occasionally drawn from a uniform distribution across the entire circle (oddball trials). Participants placed their shield on each trial (brown line) providing information about their inference about the cannon aim. **C)** In changepoint blocks, cannon aim was stationary for most trials but was occasionally resampled uniformly from possible angles (changepoint) and cannonball locations were always drawn from a Von Mises distribution centered on the true cannon aim (green points). **D&E)** Optimal inference could be approximated in both generative environments by tracking and adjusting learning according to relative uncertainty and the probability of an unlikely event (oddball or changepoint).

### Behavior of human participants and normative model

In both conditions, participants were instructed to place a shield on each trial in order to maximize the chances of blocking the upcoming cannonball (Figure 1B&C, orange line). However, behavior differed qualitatively in these two conditions, which can be observed clearly in the example participant data in Figure 1. In particular, shield placements were not updated in response to extreme outcomes in the oddball condition (oddballs; Fig 1B) but were updated dramatically in response to extreme outcomes in the changepoint condition (changepoints; Fig 1C).

To quantitatively analyze the differences between the two task conditions, we extended a previously developed normative learning model (2,7). The model approximates optimal inference using an error-driven learning rule by adjusting learning from trial to trial according to two latent variables. The first latent variable tracks the probability with which the most recent outcome was generated from an unexpected generative process (oddball probability in Fig 1D; changepoint probability in Fig 1E), whereas the second latent variable tracks the model’s uncertainty about the true cannon aim (Fig 1D&E; uncertainty). Critically, the model stipulates that surprising events in the oddball condition, which is tracked through the model’s estimate of oddball probability, should reduce learning, as oddballs are unrelated to future cannonball locations (4). In contrast, the model stipulates that surprising events in the changepoint condition, which are tracked through the model’s estimate of changepoint probability, should amplify learning, as changepoints render prior cannonballs (and thus prior beliefs) irrelevant to the problem of predicting future ones (21,22). Qualitatively, behavior from the example participant seems to follow these prescriptions, with adjustments in shield position fairly minimal on trials that include a spike in oddball probability (Fig 1B,D), but fairly large on trials that include a spike in changepoint probability (Fig 1C,E).

The normative model also makes quantitative prescriptions for how learning should be adjusted according to surprise differentially in the changepoint and oddball conditions. The surprise of a given outcome can be measured crudely through the degree to which a cannonball location differed from that which was predicted (e.g., the shield position). Larger absolute prediction errors indicate a higher degree of surprise, and higher oddball or changepoint probabilities depending on the task condition. Learning in this task can be measured through the degree to which a participant adjusts the shield position in response to a given prediction error (2), and a fixed rate of learning would correspond to a straight line mapping each prediction error onto a corresponding shield update, where the slope of the line can be thought of as the *learning rate* (Fig 2C, gray lines). The normative learning model does not prescribe a fixed learning rate across all levels of surprise; instead it prescribes higher learning rates for more surprising outcomes in the changepoint condition (Fig 2C, orange) and lower learning rates for more surprising outcomes in the oddball condition (Fig 2C, blue).

**Figure 2:**
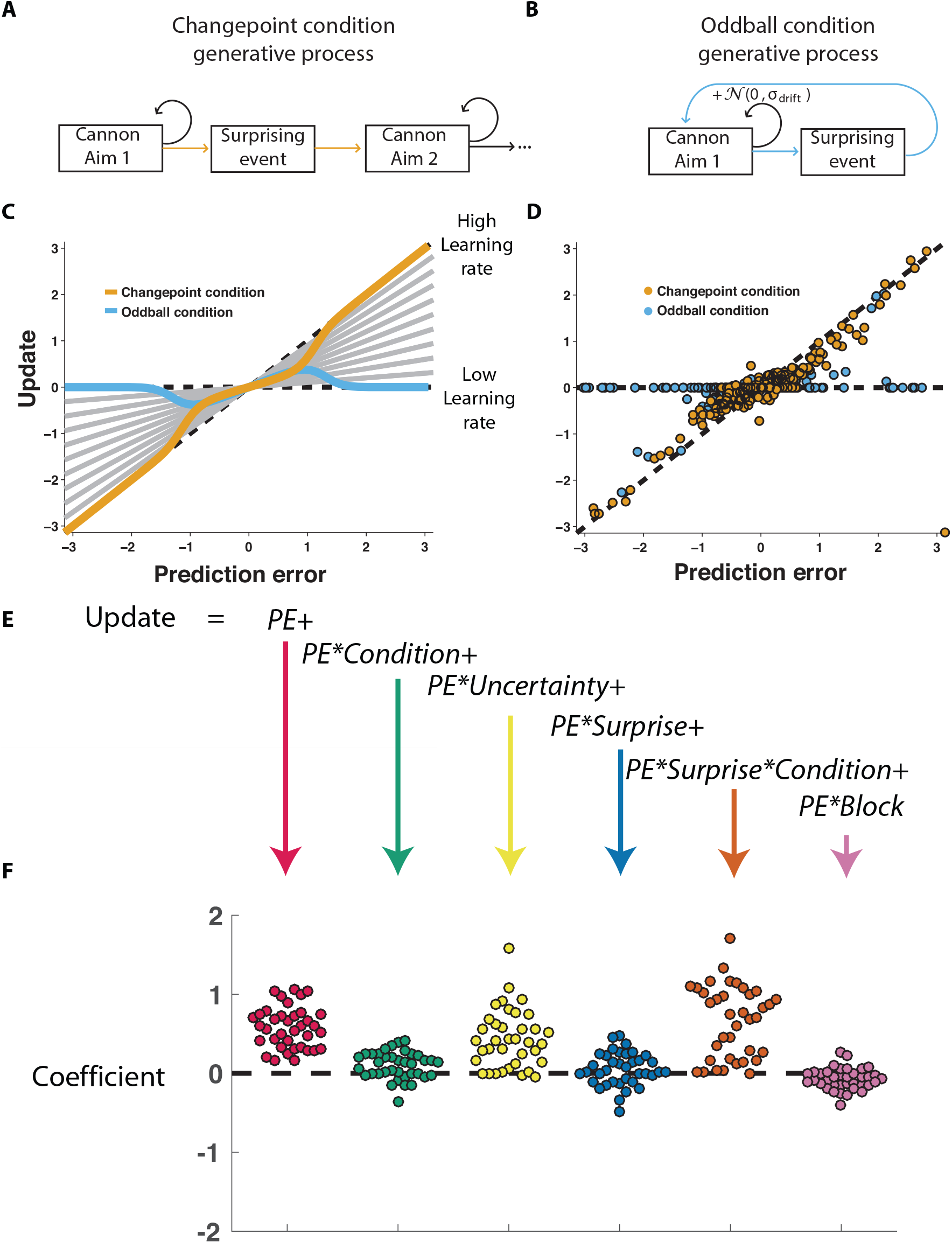
Participants scale learning according to surprise differently in changepoint and oddball contexts as would be expected for normative learning rate adjustment. **A)** In the changepoint condition, surprising events (change points) signaled a transition in the aim of the cannon whereas **B)** in the oddball condition, surprising events (oddballs) were unrelated to the process through which the aim of the cannon transitioned. **C)** Learning rate in the cannon task can be described by the slope of the relationship between prediction error (signed distance between cannonball and shield; abscissa) and update (signed change in shield position after observing new cannonball location; ordinate). Fixed learning rate updating corresponds to a line in this space whose slope is uniform across prediction errors and reflects the learning rate (gray lines). In contrast, normative learning dictates that the slope should decrease for extreme prediction errors in the oddball condition (blue) but increase for extreme prediction errors in the changepoint condition (orange). **D)** Prediction error (abscissa) and update (ordinate) for each trial (points) in each condition (designated by color) completed by a single example participant. **E)** Trial updates for each subject were fit with a regression model that included prediction errors (to measure fixed learning rate) as well as several interaction terms to assess how learning depended on various factors. **F)** Coefficients from regression model fit to individual subjects (points) revealed an overall tendency to update toward recent cannonball locations (red, t = 3.5, dof = 36, p = 10^−15^), and a tendency to do so more in the changepoint condition (green, t = 13.5, dof = 36, p = 0.001), when uncertain (yellow, t = 7.2, dof = 36, p = 2×10^−8^), and on trials where the cannonball was not blocked by the shield (pink, t = −3.3, dof = 36, p = 0.002). The model revealed that there was no consistent effect of surprise on learning across both conditions (blue, t = 1.5, dof = 36, p = 0.15), but that there was a strong interaction between surprise and condition (orange, t = 8.8, dof = 36, p = 2×10^−10^) whereby surprise tended to increase learning in the changepoint condition but decrease learning in the oddball condition.

Participants adjusted learning behavior in accordance with normative predictions, albeit with considerable heterogeneity across trials and participants. Shield updating behavior and corresponding prediction errors for an example participant reveal the basic trend predicted by the normative model, although exact updates were variable from one trial to the next (Fig 2D). To summarize the degree to which updating behavior of individual subjects was contingent on key task variables, we constructed a linear regression model that described trial-by-trial updates in terms of prediction errors as well as key task variables thought to modulate the degree to which prediction errors are translated into updates (Fig 2E) including condition (changepoint versus oddball block), surprise (as measured by changepoint or oddball probability estimates from normative model), and their multiplicative interaction (capturing the degree to which learning is increased for surprising outcomes in the changepoint context, but decreased for surprising outcomes in the oddball context). As expected, prediction error coefficients were positive, capturing a tendency for participants to update shield position toward the most recent cannonball position (Fig 1F, red; mean/SEM beta = 0.58/0.04, t = 13.5392, dof = 36, p = 10^−15^). Furthermore, participants systematically adjusted the degree to which they did so according to condition (Fig 1F, green; mean/SEM beta = 0.1/0.03, t = 3.5, dof = 36, p = 0.001), but not significantly according to surprise (Fig 1F,blue; mean/SEM beta = 0.05/0.03, t = 1.5, dof = 36, p = 0.15). Critically, surprise robustly impacted learning in opposite directions for the two conditions, as indicated by the interaction between surprise and condition (Fig 1F, orange; mean/SEM beta = 0.66/0.07, t = 8.8, dof = 36, p = 1.5 × 10^−10^). Specifically, positive coefficients indicate that sensitivity to prediction errors was increased for surprising outcomes in the changepoint condition and decreased for surprising outcomes in the oddball condition, as predicted by the normative model.

### Electrophysiological signatures of feedback processing

We took a data driven approach to identify electrophysiological signatures of feedback processing. First we regressed feedback-locked EEG data collected simultaneously with task performance onto an explanatory matrix that included separate binary variables reflecting changepoint and oddball trials, amongst other terms (Fig 3A, left). Spatiotemporal maps for changepoint and oddball coefficients were combined to create a *surprise* contrast (changepoint + oddball) and a *learning* contrast (changepoint – oddball) for each subject. Contrasts were aggregated across subjects to create a map of t-statistics (Fig 3A, right), and spatiotemporal clusters of electrode/timepoints exceeding a cluster-forming threshold were tested against a permutation distribution of cluster mass to spatially and temporally organized fluctuations in voltage that related to task variables.

**Figure 3:**
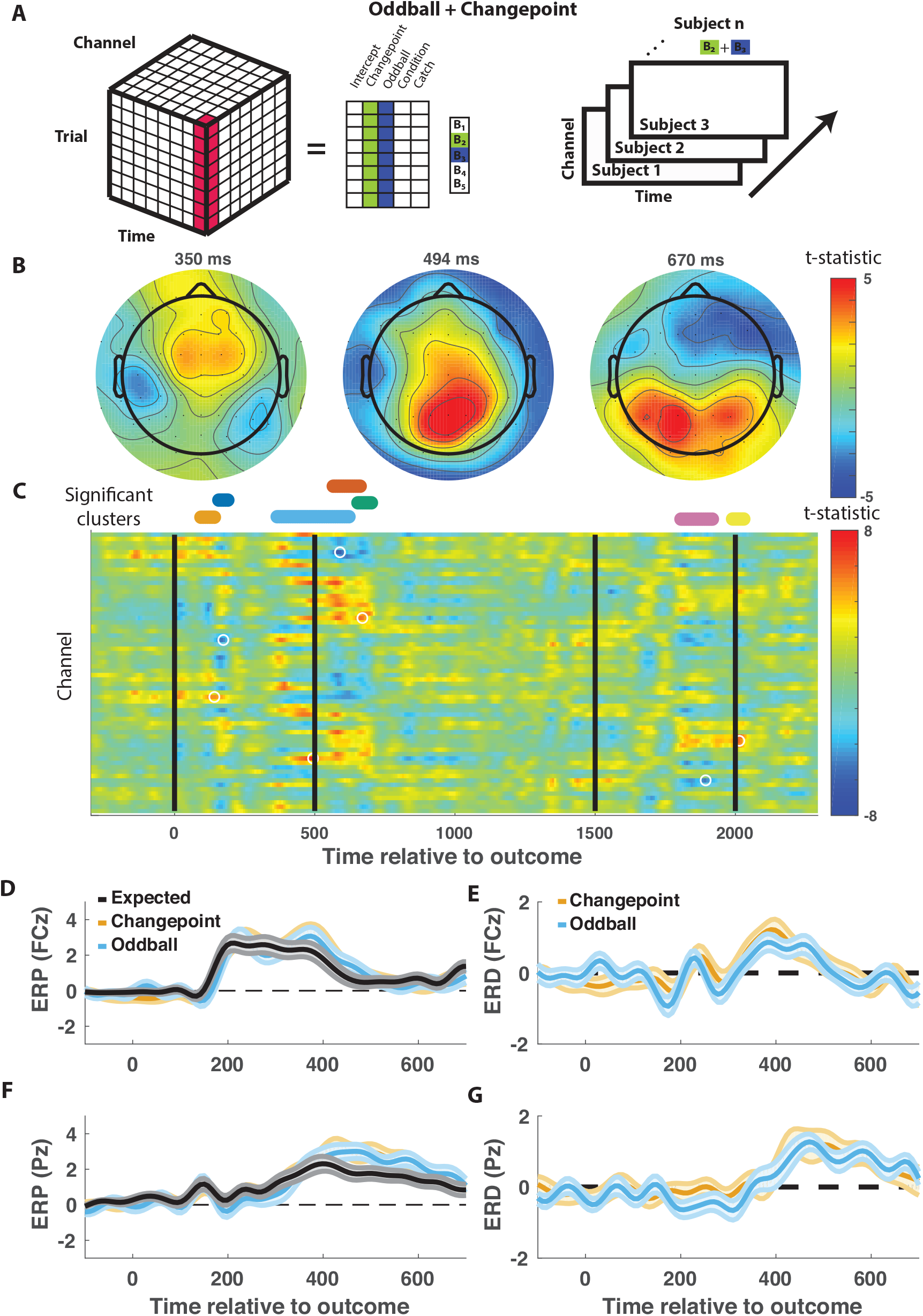
Outcome-locked central positivity reflects surprise irrespective of context. **A)** Trial-series of EEG data for a given electrode and timepoint was regressed onto an explanatory matrix that contained separate binary regressors for changepoint and oddball trials (left). A t-statisticmap was created for each electrode and time point on the surprise coefficient contrast (right). **B&C)** T-statistic map for surprise contrast across time (abscissa; **C**) and channel (ordinate; **C**) along with corresponding topoplots **B**. Separate spatiotemporal clusters that survived multiple comparisons correction via permutation testing are depicted in different colors (**C**; above heat plot). The time and channel corresponding to the maximum absolute t-statistic for each such cluster are depicted with a white circle (**C**). **D&F)** Mean/SEM (line/shading) event related potentials (microvolts) sorted by trial type (orange=changepoint, blue=oddball, black=other trials) for frontocentral (**C**; FZc) and central posterior (**F**; Pz) electrodes. **E&G**) Mean/SEM (line/shading) event related difference waveforms computed by subtracting the ERP for typical trials from the average ERP for change-point and oddball trials at frontocentral (**E**; FZc) and central posterior (**G**; Pz) electrodes.

When applied to the *surprise* contrast, this procedure yielded a large number of significant clusters distributed across electrodes and timepoints (Fig 3C). Two clusters of positive coefficients occurring 350 to 700 ms after onset of the cannonball location were of particular interest, given the consistency of their timing and direction with the canonical p300 response. Examining the spatial distribution of coefficients during this period reveals an early frontocentral locus of positive coefficients (350 ms; Fig 3B, left) that moves posterior and hits a peak t-statistic at 494 ms (Fig 3b, middle). Later on, the positive coefficients spread laterally and reach a second peak at 670ms (Fig 3B, right). The clustering procedure divides the positive coefficients observed from 350ms to 700ms into two clusters peaking at 494 and 670 ms.

The time course of positive coefficients within anterior and posterior central electrodes suggests that these clusters are picking up on P3a and P3b components of a P300 response to the outcome delivery. Average outcome-locked event related potentials in a frontocentral electrode (FCz) reveal a positive deflection from 300-500 ms (Fig 3D, black). This deflection is enhanced on both changepoint and oddball trials (Fig 3D,E, orange and blue), reminiscent of the P3a component, also referred to as the novelty P300. Posterior electrode (Pz) event-related potentials (ERPs) reveal a later and longer lasting positive deflection in response to a new outcome (Fig 3F, black). This positive deflection is enhanced on both changepoint and oddball trials (Fig 3F,G, orange and blue), reminiscent of the P3b, or updating component of the P300. Since the spatial and temporal profiles of our clusters were consistent with what has been referred to in previous literature as the P300, we will refer to the clusters peaking at 494 and 670 ms as early and late components of the P300, respectively.

In contrast to the EEG signature of *surprise*, which included a robust and extended P300 response, the only signals identified by the *learning* contrast (changepoint-oddball) were early (peak at 158 ms) and transient (Fig S3-1).

### Behavioral relevance of the P300

Competing theories posit different functional roles for the signal underlying the P300. In particular, some theories suggest that the P300 reflects a general surprise signal, whereas others attribute a more specific role in accumulating information, for example about the current state of the world. To test how early and late P300 components may relate to learning behavior in our task we extracted trial-to-trial measures of these components by taking the dot product of the cluster t-map and each single trial ERP (Fig 4A, (23)). The dot product indexes the degree to which a single trial ERP displays the profile of a given spatiotemporal cluster, thereby allowing us to test the degree to which the measured signal on any given trial might relate to behavior. We then examined how trial-to-trial behavioral updates in shield position related to these single trial EEG signal strengths using a regression model similar to that employed in the behavioral analysis (Fig 4B). The regression model included two key terms to characterize the influence of 1) the multiplicative interaction of prediction error with the EEG signal strength, and 2) the interaction between prediction error, EEG signal strength and condition. The first EEG-based term provided a measure of the relationship between learning and the P300 that was independent of condition, and thus allowed us to test the prediction that the P300 reflects a *direct learning* signal (Fig 4C). The second EEG-based term provided a measure of the relationship between learning and the P300 that depended on condition (*conditional learning*), and thus allowed us to test the prediction that any learning impact of the P300 is bidirectionally sensitive to the source of surprise (Fig 4D).

**Figure 4:**
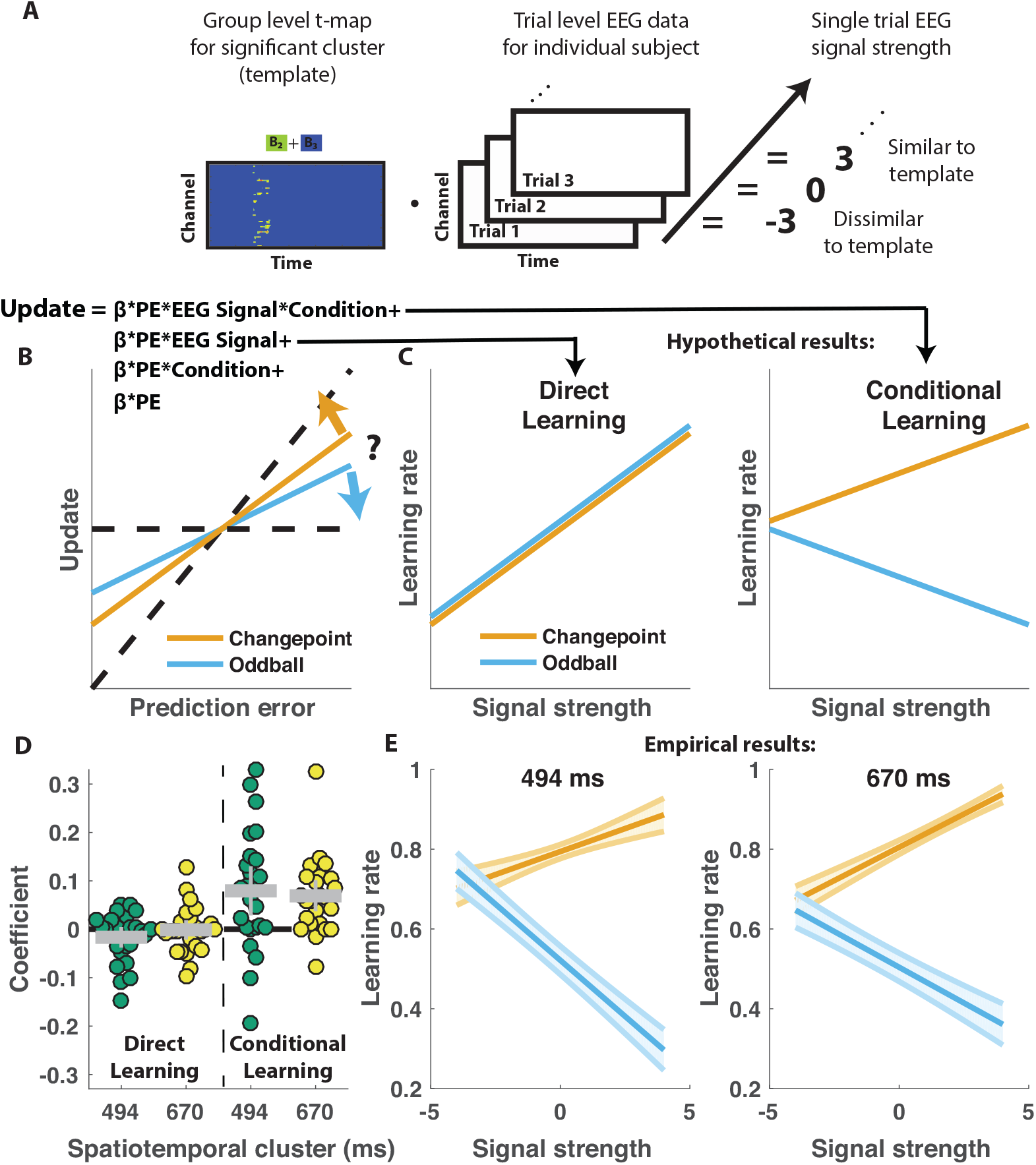
Central positivity predicts learning in opposite directions for changepoint and oddball contexts. **A)** T-maps corresponding to significant spatiotemporal clusters were used as templates to estimate trial-by-trial signal strength. **B)** Single trial updates for each participant were fit with a regression model that included additional terms to describe 1) the degree to which learning was increased on trials in which the EEG signal was stronger (PE times EEG signal) as would be expected for a canonical learning signal and 2) the degree to which learning was conditionally modulated by the EEG signal (PE times condition times EEG signal) as would be expected for a surprise signal that influenced downstream learning computations. **C)** Hypothetically, the learning rate (slope of the relationship between updates and prediction errors) might increase for stronger EEG signals (left) which would be captured by the PE times EEG direct learning regressor. Alternatively, the learning rate may increase for stronger EEG signals in the changepoint condition and decrease for stronger EEG signals in the oddball condition, as measured by the *conditional learning* regressor. **D)** Individual subject coefficients revealed no significant main effect of either early (494 ms; green) or late (674 ms; yellow) P300 signals on direct learning (left), but a strong positive interaction (*conditional learning*) effect at both time points (right), indicating that the signals were differentially predictive of learning in the changepoint and oddball conditions. **E)** Learning rates predicted by the regression model (ordinate) increased as a function of signal strength (abscissa) for each P300 cluster (left=494ms; right=674ms) in the changepoint condition (orange) but decreased as a function of signal strength in the oddball condition (blue).

Indeed, participant learning behavior systematically related to trial-by-trial measures of the P300, but only in a manner that depended critically on task condition. *Direct learning* coefficients from the model revealed that neither early (mean/SEM = −0.012/0.01, dof = 24, t = 1.6, p = 0.12) nor late (mean/SEM = −0.001/0.009, dof = 24, t = −0.12, p = 0.90) components of the P300 were systematically related to learning in the same manner across both conditions (Fig 4D, left). In contrast, *conditional learning* coefficients from both clusters tended to be positive across subjects (mean/SEM for 494,670ms cluster = 0.07/0.02, 0.069/0.02, dof = 24,24, t = 3.2, 4.5, p = 0.004, 0.0002). Learning rate predictions derived from the regression model show that higher P300 signal strength predicts more learning in the changepoint condition (Fig 4E, orange), but less learning in the oddball condition (Fig 4E, blue). Thus, there was a systematic relationship between P300 and learning, but that relationship was oppositely modulated by the task condition and hence the inferred source of surprise.

The relationship between the P300 and participant learning behavior persisted even after controlling for all known sources of variability in learning behavior. In a model of shield updating behavior that included predictions from the behavioral model described previously (Fig 2E) *conditional learning* coefficients for the late P300 component were reduced relative to the previous regression model, but still greater than zero (Fig 5B, yellow; mean/SEM for 670ms cluster = 0.03/.01 dof = 24, t = 2.5, p = 0.02). *Conditional learning* coefficients for the earlier P300 cluster were inconclusive (Fig 5B, green; mean/SEM for 494 ms cluster = 0.02/0.01, dof = 24, t = 1.6, p = 0.11). However, the participants who showed the greatest behavioral modulation of learning according to surprise and condition tended to also have the highest *conditional learning* coefficients indicating the degree to which P300 conditionally predicted learning beyond what could be achieved with our best behavioral model (Fig 5C; r = 0.42, p = 0.04). Thus, the magnitude of the P300 signal predicted learning increases in changepoint contexts and learning decreases in oddball contexts and did so beyond what could be predicted with behavioral modeling alone.

**Figure 5:**
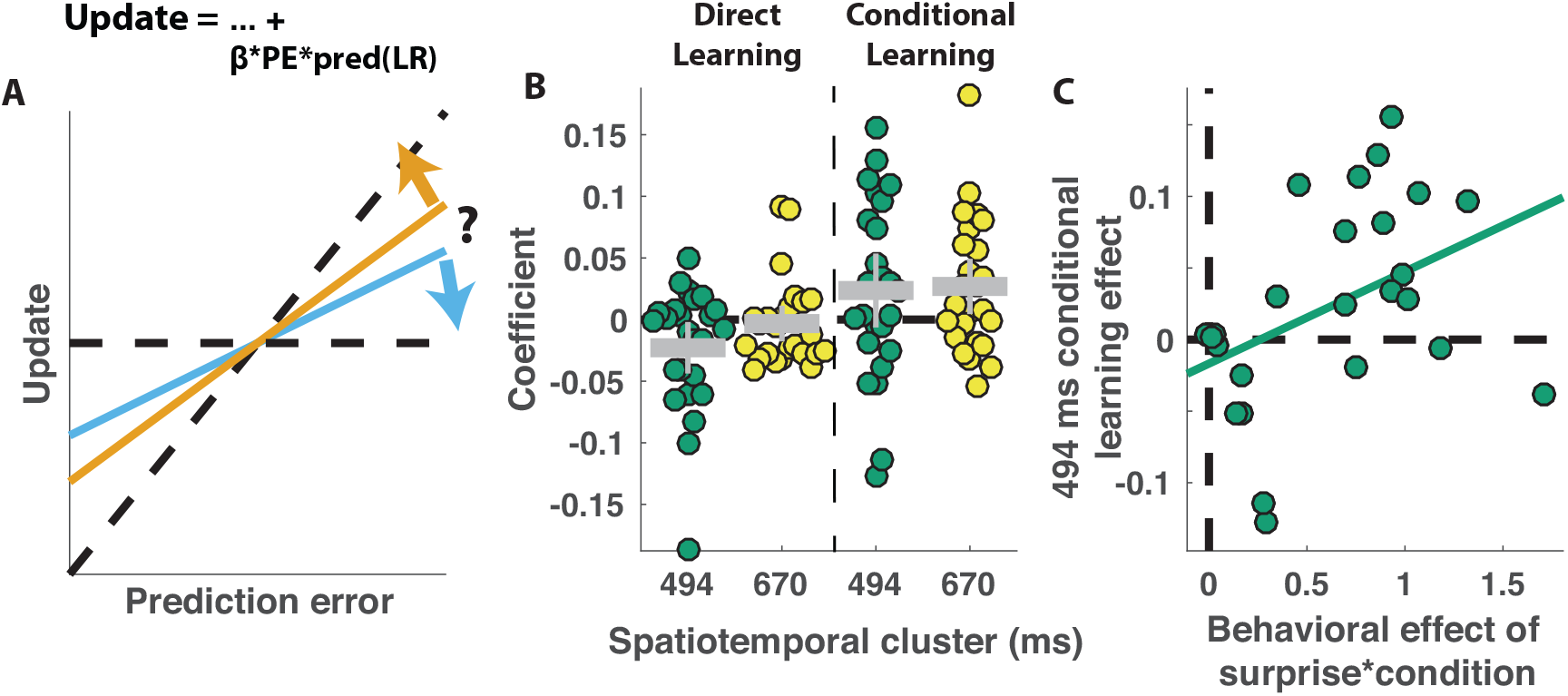
Central positivity explains trial-to-trial learning behavior that could not be otherwise captured through behavioral modeling. **A)** Single trial updates for each participant were fit with a regression model that included the best estimates of learning rate provided by our behavioral regression model (β times PE times pred(LR)) as well as additional terms to describe the degree to which learning was increased on trials in which an EEG signal was present or the degree to which learning was contextually modulated by the EEG signal. **B)** Direct learning (left) and *conditional learning* (right) coefficients for EEG terms in the regression model are plotted for early (494 ms) and late (670 ms) components of the P300 response for each subject (colored points). Gray rectangle/lines indicate group mean/SEM. *Conditional learning* coefficients were significantly greater than zero for the late P300 component (mean/SEM for 670ms cluster = 0.03/.01, dof = 24, t = 2.5, p = 0.02) indicating that this signal predicted learning in a contextual manner even after accounting for behavioral variability that could be captured by our computational model (Figure 2). **C)** *Conditional learning* coefficients for early P300 component (494 ms; ordinate) were positively related to the behavioral index of conditional surprise sensitivity (abscissa) as measured through our behavioral regression (r = 0.42, p = 0.04).

We applied the same trial-by-trial behavioral analysis to the spatiotemporal clusters identified in our learning (changepoint-oddball) contrast and did not find systematic relationships between EEG signals and learning behavior (ps for all coefficients and spatiotemporal clusters > 0.07; Fig S4-1) even when predictions from our behavioral model (Fig 2E) were not included in the analysis.

## Discussion

The brain receives a steady stream of sensory inputs, but these inputs differ dramatically from moment to moment in the degree to which they should affect ongoing inferences about the world. People and animals do not treat each datum in this stream the same, and instead tend to rely more heavily on some pieces of information than others. Identifying the mechanisms through which these adjustments occur could be an important step toward understanding why learning occurs more rapidly in some domains or for some people, yet our understanding of these mechanisms has been heavily conditioned on specific statistical contexts, namely changing environments in which the degree to which one should learn from information is closely coupled to the surprise associated with it. Here we examined how relationships between learning and a specific brain signal, the P300 evoked EEG potential, depend on the statistical context that they are measured in.

We show that the P300 relates systematically to learning, but that the direction of this relationship depends critically on the statistical context. In a context where surprising events indicated changepoints (Fig 1C,E) and participants learned more from surprising information (Fig 2), larger P300 responses predicted increased learning (Fig 4). In contrast, in a context where surprising events indicated oddballs (Fig 1B,D) and participants deemphasized surprising information (Fig 2), larger P300 responses predicted reduced learning (Fig 4). These context-dependent predictive relationships explained variance in learning beyond what could be captured through computational modeling of behavior alone (Fig 5), suggesting that the P300 signal may be involved in adjustments of learning rate, but does so by mediating the subjective response to surprise, rather than translating surprise into a conditionally appropriate learning signal.

### Implications for theories of P300 function

Our findings are consistent with a number of studies that have suggested the P300 is related to surprise (9,14,17,24), but extend them by demonstrating the role of the signal in controlling the degree to which new information affects updated beliefs. In contrast, our results are inconsistent with standard interpretations of the context updating interpretation of the P300 (17–20). If the P300 signal controlled the degree to which new information was loaded into working memory one would expect a consistent positive relationship between the P300 and learning across conditions (Fig 4C), but our results reveal that this relationship differed markedly depending on the statistical context (Fig 4F,G).

However, as is the case with many verbal theories, predictions offered by the context updating theory depend critically on the how specific concepts are linked to actual mechanistic processes. If the definition of context were changed to reflect the process that gave rise to the outcome (e.g., normal, changepoint, or oddball), for example, and we assume that participants expected each trial to be normal, then a context updating signal could account for our data (as recognizing more confident recognition of changepoints should lead to more learning, but more confident recognition of oddballs should lead to less learning). Thus, our results constrain potential interpretations of the context updating theory, although they do not falsify the theory altogether.

Similarly our results could also be viewed as constraining more recent theories about P300 signaling. One more recent theory posits that the event locked central parietal positivity reflects accumulated evidence for a particular decision or course of action (25,26). When accumulated evidence is framed in terms of the action ultimately executed (e.g., shield placement) one might extrapolate to predict that P300 would predict higher learning in both contexts, which is not what we observed (Fig 4F&G). Nonetheless, it is difficult to extrapolate decision variables to our continuous task, and there are other mechanistic schemes in which an evidence accumulation signal over a binary decision categorizing outcome type (normal versus oddball or changepoint) might give rise to our observed results. Such an explanation would also call for response inhibition to prevent premature responding before the default category (e.g., non-oddball trial) was overturned, offering a potential link to another prominent theory of P300 function (24,27). Nonetheless, our data do not arbitrate between these theories, and instead highlight their implications for learning when mechanistic interpretations are refined and applied to our task and data.

### Neural representations of surprise and updating

A key question that has motivated a number of recent studies is how does the brain represent surprise differently than the belief updating it sometimes prescribes. Under most conditions, the degree of surprise is tightly linked to the update that is required. However, recent fMRI studies have exploited cued updating paradigms (11), irrelevant stimulus dimensions (28,29), and complementary statistical contexts (4) in order to tease apart neural representations of surprise and updating. While there are trends that seem to generalize across task boundaries (for example, dorsal anterior cingulate cortex (dACC) reflecting updating in cued updating and irrelevant stimulus dimension paradigms (11,29)) there is also a good deal of inconsistency across different tasks in terms of the roles of specific signals. For example, even though BOLD responses in dACC were identified as reflecting updating in two studies, they were shown to represent surprise in another (4) and manipulations of statistical context failed to reveal *any* brain regions that provide a pure updating signal (4).

One possible explanation for this discrepancy is that the component processes of updating and non-updating might overlap in some specific paradigms. For example, the oddball outcomes that led to reduced learning in our paradigm and that of d’Acremont & Bossaerts were dissimilar to all previous outcomes and indistinguishable on other feature dimensions (in contrast to (11)). Thus, while these outcomes do not contain information pertinent to ongoing beliefs about future outcomes, they did contain information critical for perception, namely that prior expectations should not be used to bias their perceptual representations (30). Interestingly, recent work has suggested that people dynamically adjust the degree to which percepts are biased using systems, including the pupil linked arousal system, that are closely linked to the systems implicated in adjusting learning rate (6,30–34). Thus, one possible explanation for the inconsistency in previous studies attempting to dissociate surprise from updating is that these studies have differed in the degree to which they inadvertently manipulated systems for controlling perceptual biases.

Like in the previous fMRI study relying on statistical context to dissociate learning from surprise (4), our EEG results revealed a large number of signals related to surprise and no signals that convincingly reflected learning rate in a context independent manner. This comes as somewhat of a surprise given previous work identifying EEG signals analogous to a late P300 component reflecting surprise, predicting learning and influence on choice even in paradigms where this influence was unrelated to surprise (3,9,10,15,16). In line with previous work from fMRI studies, we interpret the differences in our results from what might have been predicted based on previous work as pertaining to unique strategy we employed for dissociating learning from surprise through the use of different statistical contexts.

### Mechanisms of learning rate adjustment

Our results, particularly when taken in the context of previous studies examining how the brain adjusts learning in accordance with surprise, constrain possible models of learning rate adjustment in the brain. We show that that the updating P300 signal, which positively predicts learning in changing environments (Fig 4E), also negatively predicts learning in a context with infrequent statistical outliers (Fig. 4E). Thus, in a most basic sense, our results suggest that the P300 signals reflects an early contribution to learning rate adjustment, and that this signal is untangled according to statistical context at some downstream stage of processing. The lack of robust ERP correlates of direct learning signals (Fig S3-1 & S4-1) suggests that this downstream process does not have a task-locked electrophysiological signature.

One potential mechanism for learning rate adjustment that fits well with these constraints is the notion that adjustments in learning might be implemented through flexible replacement of state representations (35–37). Learning rate adjustment is adaptive in changing environments because it can effectively partition data relevant to the current predictive context from data that are no longer relevant to prediction (21,22). One possible implementation of this partitioning would be to change the active state representations that serve as the substrate for contextual associations. Recent work has identified signals in OFC, a region implicated in representations of latent states (38), that change more rapidly during periods of rapid learning (39). If this is indeed the implementation through which learning rate adjustments occur, observed learning rate signals might actually signal the need to adjust the representation of the latent state.

Interestingly, replacement of the active latent state, or partitioning of data more generally, might also be an effective way to implement the decreased learning observed in response to surprising observations in the oddball condition of our task. In the case of an oddball, one strategy would be to recognize the oddball as having been generated by an alternative causal process (e.g., oddball distribution) and to attribute learning to a latent representation of this process (40). Under such conditions, implementation would require a surprise signal that reflects the relevance of this oddball latent state. After the new observation is attributed to the oddball context, the system would require a transition back into the original “non-oddball” state in order to make a prediction that is unaffected by the most recent oddball outcome. The more effectively surprise is recognized and responded to through state changes (e.g., the stronger the surprise signal) the more effectively this implementation would partition an oddball observation from ongoing beliefs about the standard generative process, and therefore the smaller learning rates would be. Thus, one mechanistic interpretation of the P300 results might be that it is providing a partitioning signal that results in transitions in the internal state representation, which can either increase or decrease learning depending on the statistical context.

Confirming our proposed mechanistic interpretation of these results would require future studies more closely relating P300 signals to purported state representations (39). Furthermore, given that our study relied completely on computational modeling and correlations with behavior, our results raise important questions as to whether the observed associations could be manipulated directly pharmacologically or through biofeedback paradigms. Thus, our work provides new insight into the underlying mechanisms of learning rate adjustment and the role of the P300 in this process, but leaves many unanswered questions to be addressed in future research.

## Methods

### Participants

Participants were recruited from the Brown University community: n = 37, 21 female, mean age = 20.2 (SD = 3.1, range = 18-36). Behavioral data from all participants was included in behavioral analyses. Data from 12 participants were excluded from EEG analysis due to low data quality (> 25% of epochs rejected during preprocessing). Thus, 37 participants were included in the behavioral analyses and 25 participants were included in the EEG analyses. All human subject procedures were approved by the Brown University Institutional Review Board and conducted in agreement with the Declaration of Helsinki.

### Cannon Task

Participants performed a modified predictive inference task programmed in Matlab (The Mathworks, Natick, MA), using the Psychtoolbox-2 (http://psychtoolbox.org/) package. The task was based on predictive inference tasks in which participants are asked to predict the next in a series of outcomes (2,6,7), but differed from previous such tasks the following ways: 1) the outcomes were generated from both changepoint and oddball processes to dissociate learning from surprise, 2) information necessary for performance evaluation was not available at time of outcome so that signals related to belief updating could be dissociated from valenced performance evaluation signals, 3) the task space was circular, and 4) the generative process was cast in terms of a cannon shooting cannonballs.

Participants were instructed to place a shield at some position along a circle subtending 5 degrees of visual angle in order to maximize the chances of catching a cannonball that would be shot on that trial (Fig 1a). During an instructional training period, the generative process that gave rise to cannonball locations was made explicit to participants. During this phase, participants were shown a cannon in the center of the screen. On each trial, a cannonball would be “shot” from that cannon with some angular variability (Von Mises distributed “Noise”, concentration = 10 degrees). A key manipulation in our design was how the aim of the cannon evolved from one trial to the next. The cannon would either 1) remain stationary on the majority of trials and re-aim to a random angle with an average hazard rate of 0.14 (changepoint condition) or 2) change position slightly from one trial to the next according to a Von Mises distributed random walk with mean zero and concentration 30 degrees (oddball condition). In the changepoint condition, all cannonballs were displayed as originating at the cannon in the center of the circle, whereas in the oddball condition a small fraction (0.14) of trials were oddballs, in which the cannonball location was sampled uniformly across the entire circle and the cannonball appeared without a trajectory.

After completing the instructional training, in which the generative process was fully observable, participants were asked to perform the same basic task without being able to see the cannon. In this experimental phase participants were forced to use knowledge of the generative structure gained during training, along with the sequence of prior cannonball locations, in order to infer the aim of the cannon and to inform shield placement. Participants completed four blocks of 60 trials for each task condition (changepoint and oddball) in order randomized across participants. The 240 experimental trials for each condition always followed the instructional training period for that condition in order to minimize ambiguity over which generative structure was giving rise to the experimental outcomes.

On each trial of the experimental task, participants would adjust the position of the shield through key presses (starting at the shield position from the previous trial) until they were satisfied with its location (Fig 1a; prediction phase). After participants locked in their prediction (through a key press) there was a 500 ms delay and then the cannonball location was revealed for 500 ms (Fig 1A; outcome phase). The cannonball then disappeared for 1000 ms before it reappeared, along with a full depiction of the participants shield (Fig 1A; shield phase). The shield was always centered on the position indicated by the participant during the prediction phase, but differed in size from one trial to the next in a random and unpredictable fashion that ensured subjects could not predict whether they would successfully “catch” the cannonball during the outcome phase. Thus, information provided during the outcome phase provided all necessary information to update beliefs about the cannon aim, but did not contain sufficient information to determine whether the cannonball would be successfully caught on the trial. In addition to trial feedback provided during the shield phase, participants were also provided information about their performance at the end of each block that included the fraction of cannonballs that were caught. Participants were paid an incentive bonus at task completion that was based on the number of cannonballs that were caught.

### Computational Model

Optimal inference in the changepoint condition would require considering all possible durations of stable cannon position (21,22) but can be approximated by collapsing the mixture of predictive distributions expected to arise from this optimal solution into a single Gaussian distribution, which approximates the posterior probability distribution over cannon locations, achieves near optimal inference, reduces to an error driven learning rule in which learning rate is adjusted from moment to moment according to environmental statistics, and provides a detailed account of human behavior (2,7). Similarly, the ideal observer for the oddball generative process would require tracking the predictive distributions and posterior probabilities associated with each possible sequence of oddball/non oddball trials that could have preceded the time step of interest. Like in the changepoint condition, this algorithm can be simplified by approximating the set of all possible predictive distributions with a single Gaussian distribution, leading to an error driven learning rule in which learning rate is adjusted dynamically from trial to trial, allowing us to derive normative prescriptions for learning for both conditions.

While the normative model for the changepoint condition has been described elsewhere (7) the analogous model for the oddball condition is not, and thus we describe the normative account of oddball learning in full detail. In order to minimize the differences between experienced and modeled latent variables, we formulate our model in terms of the prediction errors made by participants on each trial (rather than those that would have been made by the model) (7). On each trial of the oddball condition, the normative model: 1) updated its representation of uncertainty, 2) observed a prediction error and computed the probability that the prediction error reflects an oddball, 3) computed the normative learning rate by combining uncertainty (step 1) and oddball probability (step 2), 4) adjusted prediction about cannon position according learning rate and prediction error.

Relative uncertainty, which reflects the fraction of uncertainty about an upcoming cannonball location that is due to imperfect knowledge of the cannon aim and is analogous to the Kalman gain, was updated on each trial according to the most recent observation (which should decrease uncertainty about cannon position) and the expected drift in the aim of the cannon occurring between trials (which should increase uncertainty about cannon position). Given that relative uncertainty is expressed as a fraction of total uncertainty, it is useful to think of the numerator of the fraction, or the estimation uncertainty over possible cannon aims, which is the variance on a gaussian mixture distribution and is updated as follows:

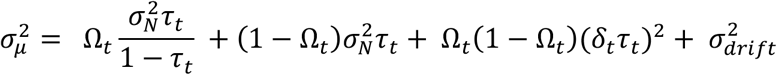

where Ω_*t*_ is the probability that an oddball occurred on trial t, 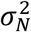 reflects the variance on the distribution of cannonball locations around the true cannon aim (noise), *τ_t_* reflects the relative uncertainty on trial t, *δ_t_* is the prediction error made in predicting the outcome on trial t, and 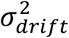 reflects the degree to which the cannon position drifts from one trial to the next. The first two terms in the model reflect the oddball and non-oddball contributions to the updated uncertainty, the third term reflects uncertainty resulting from the difference between predictions for trial t+1 conditioned on an oddball or non-oddball having occurred on trial t, and the last term reflects uncertainty resulting from the expected drift of the cannon position between trials. Relative uncertainty for trial t+1 is then updated as the updated fraction of uncertainty about the upcoming outcome that is attributable to imprecise knowledge of the true cannon position, rather than to noise in the distribution of exact cannonballs around that position:

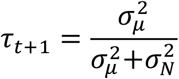

The updated relative uncertainty, along with assumed knowledge of the overall noise and hazard rate, were used to calibrate the oddball probability associated with each new prediction error:

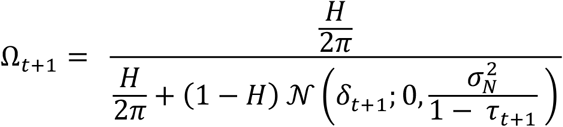

Where H is the average hazard of an oddball (0.14) and *t*_*t*+1_ is the new prediction error, and the second term in the denominator reflects the probability density on a normal distribution centered on the predicted location and with variance derived from relative uncertainty. The model’s prediction about cannon aim was then updated according to a fraction of the prediction error *t*_*t*+1_ with the exact fraction, or learning rate, determined according to the updated uncertainty and oddball probability:

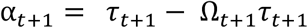

Note that relative uncertainty (*τ*_*t*+1_) contributes positively to the learning rate, whereas oddball probability (Ω_*t*+1_) reduces the learning that would otherwise be dictated by the current level of uncertainty.

### Behavioral analysis

Two key behavioral measures were extracted from each trial. First, the *prediction error* on a trial was defined as the circular distance between the cannonball location and the shield position for that trial. Second, the *update* on a given trial was defined as the circular distance between the shield position on that trial and the shield position on the subsequent trial (e.g., the updated shield position). In order to better understand the computational factors governing adjustments in shield position, we fit *updates* with a linear model that included an intercept term to model overall biases in learning along with a prediction error term to capture general tendencies to adjust the shield towards the most recent cannonball location. The model also included additional terms to model how the influence of recent cannonball locations changed dynamically according to task context. These terms included: 1) prediction error times uncertainty interaction (to model how much more participants updated shield position under conditions of uncertainty – as assessed by the computational model), 2) prediction error times surprise (where surprise was indexed by changepoint probability or oddball probability from computational model depending on the context), 3) prediction error times surprise times condition (where condition was +1 for changepoint blocks and −1 for oddball conditions), 4) prediction error times block (a categorical variable indicating whether the shield “blocked” the most recent cannonball. The regression model was fit to each participant and t-tests were performed on the regression coefficients across participants to test for significant contributions of each term to update behavior.

### EEG Acquisition

EEG was recorded from a 64-channel Synamps2 system (0.1–100 Hz bandpass; 500 Hz sampling rate). Continuous EEG data was epoched with respect to the outcome presentation for each trial. Preprocessing was done manually in Matlab (Mathworks, Natick MA) using the EEGLAB toolbox (https://sccn.ucsd.edu/eeglab/index.php) as described previously (23) and included the following steps: 1) epoching and alignment to outcome onset, 2) epoch rejection by inspection, 3) channel removal and interpolation by inspection, 4) bandpass filtering [.5-50 hz], 5) removal of blink and eye movement components using ICA. Participants for whom more than 25 percent of epochs were rejected were not included in analyses of EEG data.

### EEG Analysis

EEG Data for individual participants were analyzed using a mass univariate approach. Specifically, the trial series EEG data for a given participant, channel, and time relative to outcome onset was regressed onto an explanatory matrix that included the following explanatory variables: 1) intercept, 2) changepoint, 3) oddball, 4) condition, 5) catch. Explanatory variables 2 & 3 were binary variables marking trials in which a surprising event occurred (i.e. changepoint or oddball) whereas 4 reflected the overall task context (i.e. whether oddballs or changepoints were present in the current statistical context), and 5 conveyed whether the participant successfully “caught” the cannonball on each trial. Surprise and learning contrasts were created as the sum and difference of the changepoint and oddball coefficients, respectively. T-statistics were computed across subjects to assess the consistency of contrasts at each electrode and timepoint.

T-statistic maps were thresholded (cluster forming threshold of p=0.001, 2 tailed) and spatiotemporal clusters were identified as temporally and/or spatially contiguous signals that shared a common sign of effect and exceeded the cluster-forming threshold. Cluster mass was computed as the average absolute t-statistic within a cluster times its size (number of electrode timepoints contained within it). Cluster mass for each spatiotemporal cluster was compared to a permutation distribution for cluster mass generated using sign flipping to correct for multiple comparisons (41).

Trial-to-trial EEG analyses were conducted by computing the dot product of the t-statistic map for a given spatiotemporal cluster and the ERP measured on a given trial. The resulting measure of EEG signal strength was then z-scored across all trials and included in a behavioral regression model to explain trial-to-trial updating behavior. Like for the behavioral analyses, trial-to-trial updates were regressed onto an explanatory matrix that included intercept and prediction error terms to capture updating biases and static tendencies to update toward recent cannonball locations. In addition, EEG informed regression models included 1) the interaction between the EEG signal strength computed above and prediction error (direct learning), and 2) the three-way interaction between EEG signal strength, prediction error, and condition (conditional learning). Positive *direct learning* coefficients indicated an unconditional increase in learning for trials in which EEG signal strength was greater, whereas positive conditional learning coefficients indicated a positive relationship between EEG signal strength and learning in the changepoint condition but a negative relationship between EEG signal strength in the oddball condition. In order to test the degree to which EEG-updating relationships persisted after accounting for variability in behavior that could be captured by our computational model, we also used a version of the EEG informed regression that additionally included the predicted update from the behavioral model (y-hat) as an explanatory variable (Fig 5).

---

**Figure S3-1:**
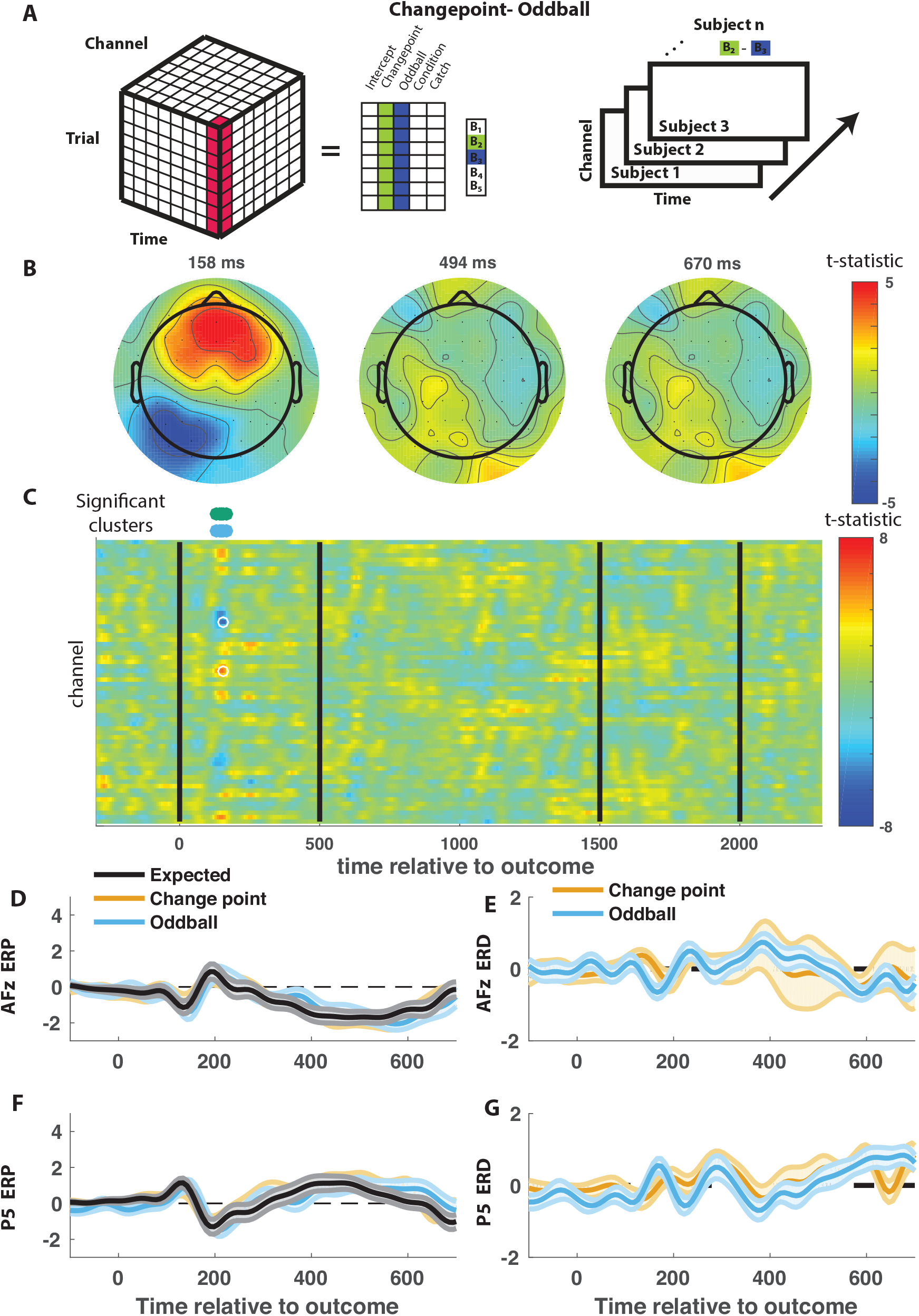
Early frontal, but not late central, positivity is greater in changepoint than oddball trials. **A)** Trial-series of EEG data for a given electrode and timepoint was regressed onto an explanatory matrix that contained separate binary regressors for changepoint and oddball trials (left). A t-statistic map was created for each electrode and time point on the learning contrast (right). **B&C)** T-statistic map for learning contrast across time (abscissa; **C**) and channel (ordinate; **C**) along with corresponding topoplots (**B**). Separate spatiotemporal clusters that survived multiple comparisons correction via permutation testing are depicted in different colors (**C**; above heat plot). **D&F)** Mean/SEM (line/shading) event related potentials (microvolts) sorted by trial type (orange=changepoint, blue=oddball, black=other trials) for frontal (**C**; AFz) and posterior (**F**; P5) electrodes that distinguish between changepoints and oddballs. **E&G)** Mean/SEM (line/shading) event related difference waveforms computed by subtracting the ERP for typical trials from the average ERP for change-point and oddball trials at frontocentral (**E**; AFz) and central posterior (**G**; P5) electrodes.

**Figure S4-1:**
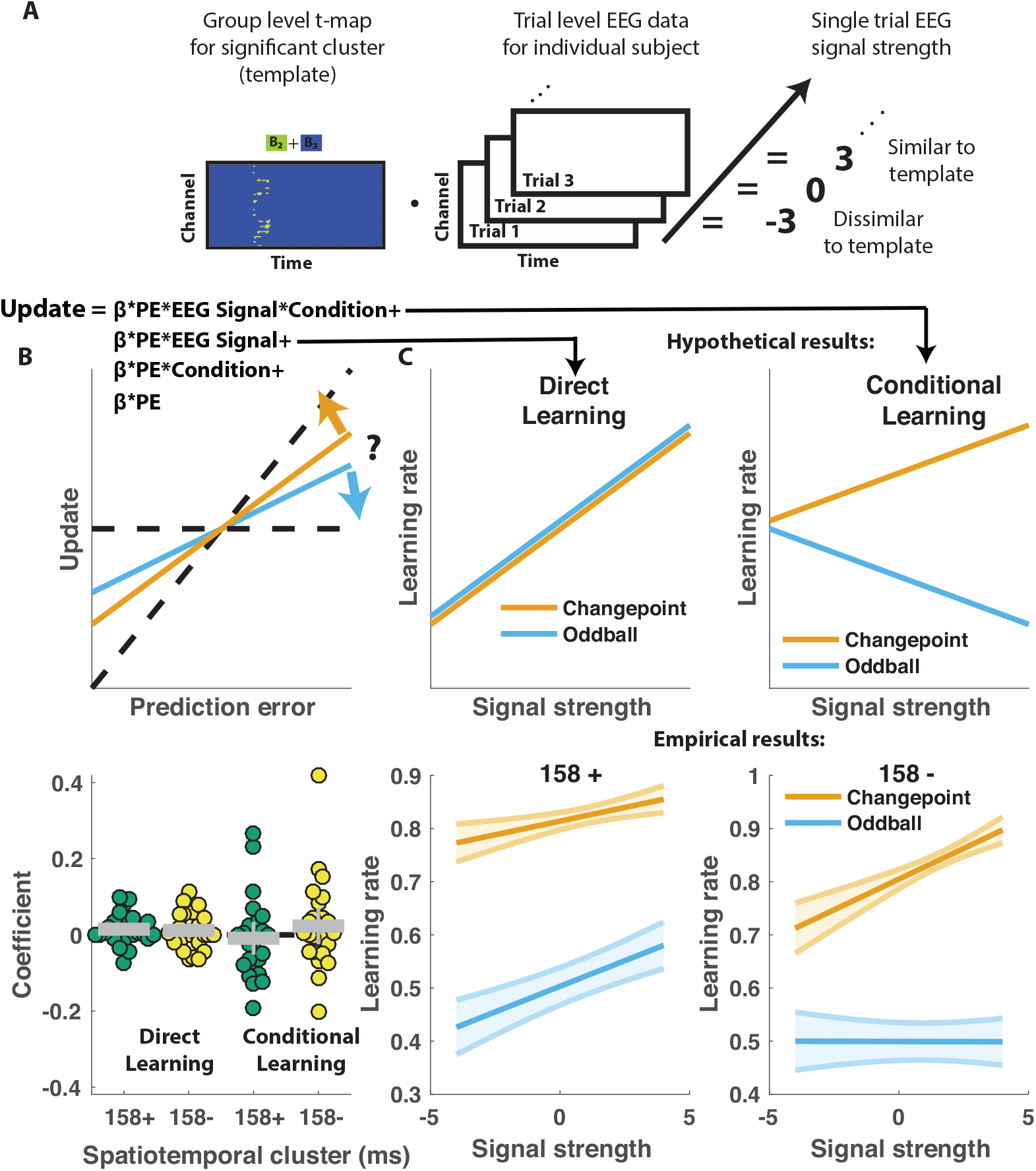
Weak direct relationships between EEG signals and learning. **A)** T-maps corresponding to significant spatiotemporal clusters were used as templates to estimate trial-by-trial signal strength (as the cosine product of the template with the outcome locked EEG data recorded on each trial). **B)** Single trial updates for each participant were fit with a regression model that included additional terms to describe 1) the degree to which learning was increased on trials in which the EEG signal was stronger (PE times EEG signal) as would be expected for a canonical learning signal and 2) the degree to which learning was contextually modulated by the EEG signal (PE times condition times EEG signal) as would be expected for a surprise signal that influenced downstream learning computations. **C)** Hypothetically, the learning rate (slope of the relationship between updates and prediction errors) might increase for stronger EEG signals (left) which would be captured by the (PE times EEG signal) regressor (*direct learning*). Alternatively, the learning rate may increase for stronger EEG signals in the changepoint condition and decrease for stronger EEG signals in the oddball condition, consistent with *conditional learning* signal. **D)** Individual subject coefficients from the changepoint-oddball spatiotemporal clusters (figure S3-1) revealed no systematic *direct learning* effect of either positive (158 +; green) or negative (158 -; yellow) EEG signals (left) (mean[SEM] coefficient = 0.01[0.008], 0.01[0.01], p = 0.08, 0.25). Nor did coefficients reveal *conditional learning* effect (right) (mean[SEM] coefficient = −0.01[0.02], 0.02[0.02], p = 0.67, 0.32). **E)** Learning rates predicted by the regression model (ordinate) tended to increase as a function of signal strength (abscissa) for both clusters (left=158+; right=158-) in the changepoint condition (orange) but were less consistent in the oddball condition (blue).

## Acknowledgements

We would like to thank Julie Helmers and Andrea Mueller for their help collecting EEG and behavioral data. This work was funded by NIH grants F32MH102009 and K99AG054732 (MRN), NIMH R01 MH080066-01 and NSF Proposal #1460604 (MJF). RB was supported by a Promos travel grant from the German Academic Exchange Service (DAAD). The funders had no role in study design, data collection and analysis, decision to publish or preparation of the manuscript.

## References

1. Behrens TEJ, Woolrich MW, Walton ME, Rushworth MFS. Learning the value of information in an uncertain world. Nature Neuroscience. 2007 Sep;10(9):1214–21.

2. Nassar MR, Wilson RC, Heasly B, Gold JI. An approximately Bayesian delta-rule model explains the dynamics of belief updating in a changing environment. Journal of Neuroscience. 2010 Sep 15;30(37):12366–78.

3. Cheadle S, Wyart V, Tsetsos K, Myers N, de Gardelle V, Castañón SH, et al. Adaptive Gain Controlduring Human Perceptual Choice. Neuron. Elsevier Inc; 2014 Mar 19;81(6):1429–41.

4. d’Acremont M, Bossaerts P. Neural Mechanisms Behind Identification of Leptokurtic Noise and Adaptive Behavioral Response. Cerebral Cortex. 2016 Feb 4;:bhw013.

5. Diederen KMJ, Spencer T, Vestergaard MD, Fletcher PC, Schultz W. Adaptive Prediction Error Coding in the Human Midbrain and Striatum Facilitates Behavioral Adaptation and Learning Efficiency. Neuron. Elsevier Inc; 2016 Jun 1;90(5):1127–38.

6. Nassar MR, Rumsey KM, Wilson RC, Parikh K, Heasly B, Gold JI. Rational regulation of learning dynamics by pupil-linked arousal systems. Nature Neuroscience. 2012 Jul;15(7):1040–6.

7. Nassar MR, Bruckner R, Gold JI, Li S-C, Heekeren HR, Eppinger B. Age differences in learning emerge from an insufficient representation of uncertainty in older adults.Nature Communications. 2016;7:11609.

8. McGuire JT, Nassar MR, Gold JI, Kable JW. Functionally dissociable influences on learning rate in a dynamic environment. Neuron. 2014 Nov 19;84(4):870–81.

9. Jepma M, Murphy PR, Nassar MR, Rangel-Gomez M, Meeter M, Nieuwenhuis S. Catecholaminergic Regulation of Learning Rate in a Dynamic Environment. O’Reilly JX, editor. PLoS Comput Biol. 2016 Oct 28;12(10):e1005171.

10. Jepma M, Brown SBRE, Murphy PR, Koelewijn SC, de Vries B, van den Maagdenberg AM, et al. Noradrenergic and Cholinergic Modulation of Belief Updating. J Cogn Neurosci. 2018 Jul 31;:1–18.

11. O’Reilly JX, Schüffelgen U, Cuell SF, Behrens TEJ, Mars RB, Rushworth MFS. Dissociable effects of surprise and model update in parietal and anterior cingulate cortex. Proceedings of the …. 2013.

12. Iglesias S, Mathys C, Brodersen KH, Kasper L, Piccirelli M, Ouden den HEM, et al. Hierarchical Prediction Errors in Midbrain and Basal Forebrain during Sensory Learning. Neuron. Elsevier Inc; 2013 Oct 16;80(2):519–30.

13. Summerfield C, Tsetsos K. Do humans make good decisions? Trends in Cognitive Sciences. Elsevier Ltd; 2015 Jan 1;19(1):27–34.

14. Garrido MI, Teng CLJ, Taylor JA, Rowe EG, Mattingley JB. Surprise responses in the human brain demonstrate statistical learning under high concurrent cognitive demand. npj Science Learn. 2016 Jun 8;1(1):e1002999.

15. Fischer AG, Ullsperger M. Real and Fictive Outcomes Are Processed Differently but Converge on a Common Adaptive Mechanism. Neuron. 2013 Sep;79(6): 1243–55.

16. Wyart V, de Gardelle V, Scholl J, Summerfield C. Rhythmic Fluctuations in Evidence Accumulation during Decision Making in the Human Brain. Neuron. Elsevier Inc; 2012 Nov 21;76(4):847–58.

17. Donchin E. Presidential address, 1980. Surprise!…Surprise? Psychophysiology. 1981 Sep;18(5):493–513.

18. Donchin E, Coles MGH. Is the P300 component a manifestation of context updating? Behav Brain Sci. 2010 Feb 4;11(03):357.

19. Polich J. THEORETICAL OVERVIEW OF P3a AND P3b. 2003 Jan 24;:1–2.

20. Polich J. Updating P300: An integrative theory of P3a and P3b. Clinical Neurophysiology. 2007 Oct;118(10):2128–48.

21. Prescott Adams R, MacKay DJC. Bayesian Online Changepoint Detection. eprint arXiv:07103742. 2007 Oct 1;:-.

22. Wilson RC, Nassar MR, Gold JI. Bayesian online learning of the hazard rate in change-point problems. Neural Comput. 2010 Sep 1;22(9):2452–76.

23. Collins AGE, Frank MJ. Within-and across-trial dynamics of human EEG reveal cooperative interplay between reinforcement learning and working memory. Proceedings of the National Academy of Sciences. 2018 Mar 6;115(10):2502–7.

24. Wessel JR. A Neural Mechanism for Surprise-related Interruptions of Visuospatial Working Memory. Cerebral Cortex. 2016 Dec 1;28(1):199– 212.

25. Kelly SP, O’Connell RG. Internal and external influences on the rate of sensory evidence accumulation in the human brain. Journal of Neuroscience. 2013 Dec 11;33(50):19434–41.

26. O’Connell RG, Dockree PM, Kelly SP. A supramodal accumulation-to-bound signal that determines perceptual decisions in humans. Nature Neuroscience. 2012 Oct 28;15(12):1729–35.

27. Wessel JR, Aron AR. Perspective. Neuron. Elsevier Inc; 2017 Jan 18;93(2):259–80.

28. Schwartenbeck P, FitzGerald THB, Dolan R. Neural signals encoding shifts in beliefs. NeuroImage. The Authors; 2016 Jan 15;125(C):578–86.

29. Nour MM, Dahoun T, Schwartenbeck P, Adams RA, FitzGerald THB, Coello C, et al. Dopaminergic basis for signaling belief updates, but not surprise, and the link to paranoia. Proceedings of the National Academy of Sciences. 2018 Oct 23;115(43):E10167–76.

30. Krishnamurthy K, Nassar MR, Sarode S, Gold JI. Arousal-related adjustments of perceptual biases optimize perception in dynamic environments. Nat hum behav. 2017 May 8;1:0107.

31. Nieuwenhuis S, De Geus EJ, Aston-Jones G. The anatomical and functional relationship between the P3 and autonomic components of the orienting response. Psychophysiology. 2011 Jan 6;48(2):162–75.

32. Vazey EM, Moorman DE, Aston-Jones G. Phasic locus coeruleus activity regulates cortical encoding of salience information. Proceedings of the National Academy of Sciences. 2018 Oct 2;115(40):E9439–48.

33. Urai AE, Braun A, Donner TH. Pupil-linked arousal is driven by decision uncertainty and alters serial choice bias. Nature Communications. 2017 Mar 3;8:14637.

34. de Gee JW, Colizoli O, Kloosterman NA, Knapen T, Nieuwenhuis S, Donner TH. Dynamic modulation of decision biases by brainstem arousal systems. eLife. 2017 Apr 11;6.

35. Collins A, Koechlin E. Reasoning, Learning, and Creativity: Frontal Lobe Function and Human Decision-Making. O’Doherty JP, editor. PLoS Biol. 2012 Mar 27;10(3):e1001293.

36. Collins AGE, Frank MJ. Cognitive control over learning: creating, clustering, and generalizing task-set structure. Psychological Review. 2013 Jan;120(1):190–229.

37. Wilson RC, Takahashi YK, Schoenbaum G, Niv Y. Orbitofrontal cortex as a cognitive map of task space. Neuron. 2014 Jan 22;81(2):267–79.

38. Schuck NW, Cai MB, Wilson RC, Niv Y. Human Orbitofrontal Cortex Represents a Cognitive Map of State Space. Neuron. Elsevier Inc; 2016 Sep 21;91(6):1402–12.

39. Nassar MR, McGuire JT, Ritz H, Kable JW. Dissociable Forms of Uncertainty-Driven Representational Change Across the Human Brain. Journal of Neuroscience. 2019 Feb 27;39(9):1688–98.

40. Gershman SJ, Niv Y. Learning latent structure: carving nature at its joints. Current Opinion in Neurobiology. 2010 Apr;20(2):251–6.

41. Nichols TE, Holmes AP. Nonparametric permutation tests for functional neuroimaging: a primer with examples. Hum Brain Mapp. 2002 Jan;15(1):1–25.

